# Mapping the spatial proteomic signature of dorsal and ventral hippocampus in a mouse model of early Alzheimer’s disease: changes in synaptic plasticity-related proteins associated with sexual dimorphism

**DOI:** 10.1101/2024.05.08.593134

**Authors:** Ana Contreras, Raquel Jiménez-Herrera, Souhail Djebari, Juan D. Navarro-López, Lydia Jiménez-Díaz

## Abstract

**Background:** An initial neuropathological hallmark of Alzheimer’s disease (AD) is the hippocampal dysfunction caused by amyloid-*β* (A*β*) peptides accumulation. Soluble oligomeric forms of A*β* shift synaptic plasticity induction threshold leading to memory deficits in male and female mice in early amyloidosis models. Some protein changes underlying those deficits have been previously studied, but the spatial distribution within the hippocampus, as well as the potential sex differences, remain unknown. Since each hippocampal region (dorsal *vs*. ventral) has clearly distinct functionality and connectivity, we postulated that some protein changes may be unique to each and might also be sex-dependent.

**Methods:** An innovative spatial proteomics study was performed to map whole hippocampal proteome distribution using matrix-assisted laser desorption/ionization (MALDI) imaging mass spectrometry, which allows protein detection with spatial resolution directly on tissue sections. Brains from sixteen adult male and female mice intracerebroventricularly injected with A*β*_1-42_ oligomers or vehicle were sectioned. MALDI imaging was performed using a RapifleXTM MALDI TissuetyperTM TOF/TOF mass spectrometer followed by protein identification by traditional tandem mass spectrometry (MS/MS) directly on the tissue. To precisely delineate both dorsal and ventral hippocampus, a Nissl staining was performed on succeeding tissue sections.

**Results:** Of the 234 detected peptides, significant differences in expression levels were found in 34 proteins, due to treatment, sex, or hippocampal location. Moreover, a significant protein-protein interaction (PPI) was observed, showing a relationship to long-term potentiation (LTP), the functional basis of memory. Accordingly, 14 proteins related to synaptic plasticity and/or AD were selected to further study. Results showed many of the altered protein to modulate glycogen synthase kinase-3*β* (GSK-3*β*), a protein widely involved in the regulation of synaptic plasticity induction threshold. In fact, hippocampal GSK-3*β* was found overactivated suggesting a facilitated long-term depression (LTD) instead of LTP in AD models.

**Conclusions:** This study offers for the first time the specific protein changes in dorsal/ventral hippocampus in both male and female mice, that modulate GSK-3*β* activity, providing new insight in the pathogenesis of early AD and valuable potential biomarkers for early diagnosis and therapeutic targets.

## 1. Background

Alzheimer’s disease (AD) is the most prevalent cause of dementia, accounting for an estimated 60% to 80% of the 55 million cases of dementia worldwide [1]. One of its early hallmarks is the presence of amyloid-*β* (A*β*) peptides [2]. The hippocampus, a critical brain area related to learning and memory, is one of the first regions affected by early amyloidosis through impairing excitatory/inhibitory (E/I) neurotransmission balance, synaptic plasticity and oscillatory activity, together leading to learning and memory disfunction [3–5].

Amyloidogenic processing of amyloid precursor protein (APP) causes the aggregation of different A*β* species, being the 42 amino acid-long amyloid-*β* (A*β*_1-42_) the dominant form in amyloid plaques in AD patients [6, 7]. However, the physiopathology of AD starts decades before amyloid plaques are present, when A*β* is soluble rather than accumulated [8]. A*β*_1-42_ monomers are highly prone to aggregation, and they form a wide range of soluble oligomers, from dimers to trimers, and ultimately fibrillate to form A*β* plaques [9, 10]. Soluble A*β*_1-42_ oligomers (oA*β*_1-42_) are the major toxic agents in early AD, leading to initial loss of excitatory synapses and blockage of synaptic plasticity both *in vivo* and *in vitro* [11–13].

To study those early stages of AD, our group and others have validated a murine model of intracerebroventricular (*icv.*) oA*β*_1-42_ administration, that shows E/I imbalance, deficits in long-term synaptic potentiation (LTP) induction, disruption of neural oscillatory synchronization and impairments in learning and memory, in both male and female mice [14–18].

Several omics techniques have been widely used to obtain an unbiased insight into the molecular changes in the progression of AD, such as genomics [19], proteomics [20, 21] and metabolomics [22]. In fact, a specific plasma proteomic profiling has shown AD stage-dependent dysregulations [23], highlighting the importance of biomarker discovery for early identification of AD risk factors based on the proteomic profile. However, most of them are carried out in cerebrospinal fluid [24], plasma [23] or brain homogenates [25–28], lacking the information on the spatial distribution of those potential biomarkers in the brain. Spatial information is key, since first A*β* deposits and subsequently tau tangles brain distribution correlate with the regional atrophy and cognitive decline at different AD stages [29–32]. In this regard, matrix-assisted laser desorption/ionization (MALDI) imaging mass spectrometry has arisen in the last decade as a particularly powerful emerging technology, that provides molecular mass determination of analytes directly on the tissue, with spatial resolution. Using MALDI imaging, previous works have revealed differential distribution of lipids, glycomes and metabolites in different brain areas both in AD patients and murine models [33–36]. A recent study has shown alterations in proteins related to synaptic function and neurodegeneration, specifically in the cortex, the ventricular zone and the corpus callosum in neonatal 5xFAD mice using MALDI imaging [37]. Nevertheless, the differential distribution of proteins due to early amyloidosis impairment in the hippocampus has not been studied, which is particularly important due to its functional, molecular and connectivity heterogeneity [38–40].

Here, for the first time, the hippocampal proteome of healthy and early amyloidosis treated male and female mice was investigated, and potential protein expression differences between dorsal and ventral hippocampal areas were analyzed, providing cutting-edge data on the specialization of the hippocampal proteome and valuable potential biomarkers for early diagnosis and therapeutic targets.

## 2. Methods

### 2.1. Animals

Eight female and eight male C57BL/6 adult mice (12-24 weeks old; 20-30 g) were used (RRID:MGI:5,656,552; Charles River, USA). Animals were kept on 12 h light/dark cycles with access to food and water *ad libitum* and controlled temperature (21 ± 1°C) and humidity (50 ± 7%). Mice were housed in same-sex groups of 5 per cage before surgery, and individually afterwards. Environmental enrichment elements were provided. All experimental procedures were carried out at the same time interval in both female and male mice to minimize circadian rhythm interferences.

All experimental procedures were reviewed and approved by the Ethical Committee for Use of Laboratory Animals of the University of Castilla-La Mancha (PR-2021-12-21) and conducted according to the European Union guidelines (2010/63/EU) and the Spanish regulations for the use of laboratory animals in chronic experiments (RD 53/2013 on the care of experimental animals: BOE 08/02/2013).

### 2.2. Surgery for early hippocampal amyloidosis model generation

For anesthesia induction, 4% isoflurane (#13400264, ISOFLO, Proyma S.L., Spain) was administered using a calibrated R580S vaporizer (RWD Life Science; flow rate: 0.5 L/min O_2_). During the remaining part of the procedure, 1.5% isoflurane was delivered constantly for anesthesia maintenance, after which intramuscular buprenorphine (0.01mg/kg; #062009, BUPRENODALE, Albet, Spain) and a healing cream (Blastoestimulina; Almirall, Spain) were administered to accelerate recovery and decrease animal suffering.

Mice were implanted with a blunted, stainless steel, 26-G guide cannula (Plastics One, US) in the left ventricle (1 mm lateral and 0.5 mm posterior to bregma; depth from brain surface, 1.8 mm) [41] for *icv*. administration of A*β.* The final position of the cannula was determined by Nissl staining after brain tissue collection [18].

As previously described [42], A*β*_1-42_ (#AB120301; Abcam, UK) was dissolved in phosphate-buffered saline (PBS) and incubated 6 h at 37°C to form soluble oA*β*_1-42_ before administration [43]. After a week recovery time at its minimum, freely moving animals received a 3 μL *icv.* injection of either 1 μg/μL of oA*β*_1-42_ or vehicle (PBS), through a 33-G internal cannula within the implanted guide cannula and protruding 0.5 mm into the ventricle, using a motorized Hamilton syringe at a rate of 0.5 μL/min (Figure 1A). After administration, the internal cannula was not removed for an extra minute to avoid backflow. The dose was selected based on previous studies that demonstrated its effectiveness and safety [14–16, 18].

**Figure 1.**
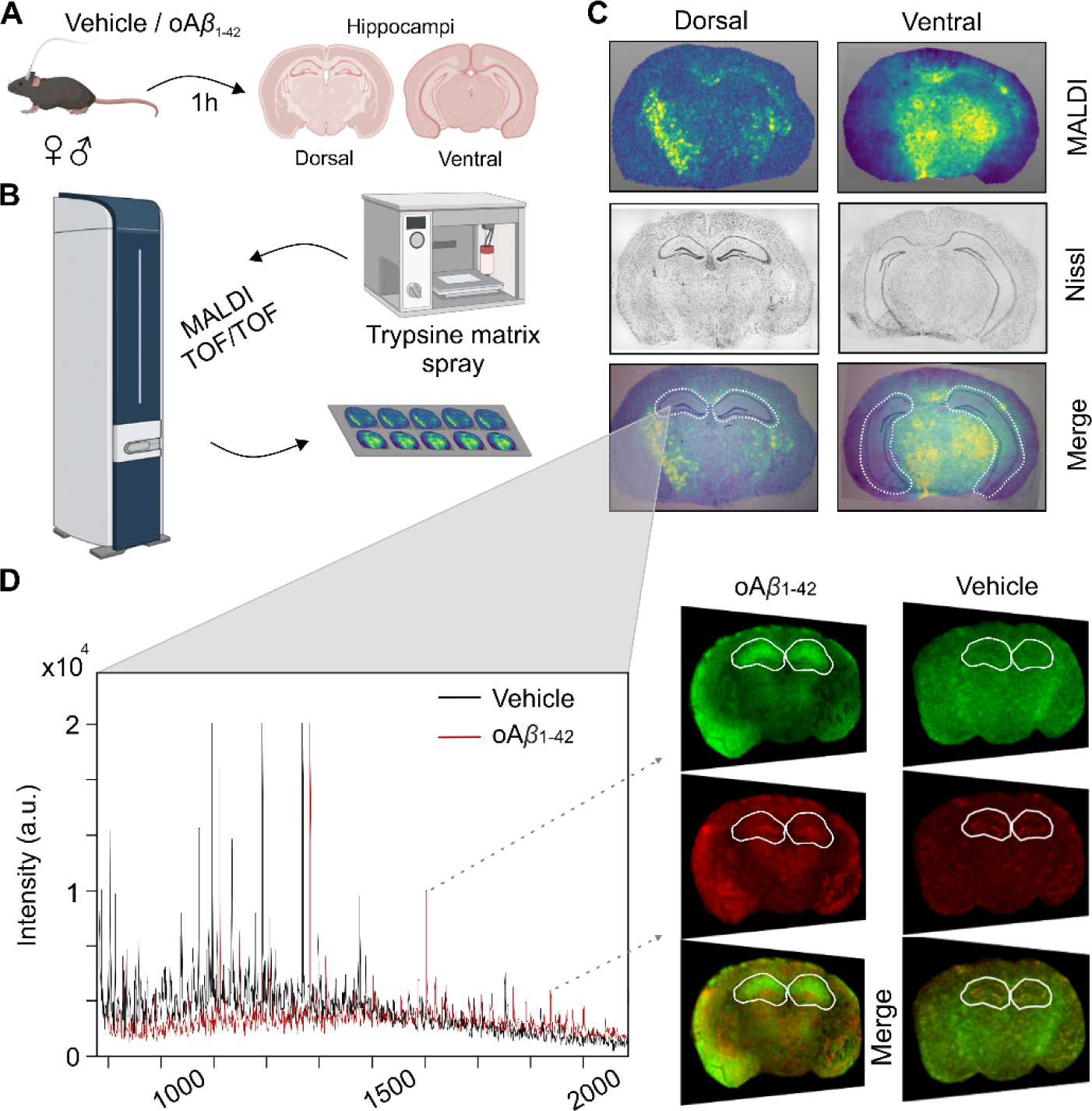
Experimental design. **(A)** Schematic illustration of *icv*. injection of either vehicle or oA*β*_1-42_ in male and female mice and dissection of dorsal- and ventral- containing hippocampal sections1 hour later. **(B)** Schematic representation of trypsin and CHCA matrix spray application with a HTX TM SprayerTM and subsequent MALDI imaging acquisition using a RapifleXTM MALDI TissuetyperTM TOF/TOF mass spectrometer. **(C)** MALDI images showing protein markers (top), Nissl staining of consecutive tissue sections (middle, in gray) and overlay of the two (bottom), with delimitation of dorsal (left) and ventral (right) hippocampi in white (dashed line). **(D)** Representative peptide spectrum (left) of hippocampal tissue measured by MALDI imaging from vehicle (black) and oA*β*_1-42_-treated (red) mice with examples (right) of two selected peptides shown in tissue sections (green and red) and its merge.

### 2.3. Brain tissue collection and preparation for MALDI MS imaging

One hour after treatment, animals were decapitated, and brain was extracted and dissected (Figure 1A). This time point was chosen based on previous experiments showing that even 1 h post-*icv*. injection spatial memory impairment was already evident in our animal model of early acute hippocampal amyloidosis [18]. The medial part of the brain, containing the hippocampus, was fresh frozen at −80°C until further use. Hippocampal-containing sections were cut into 10 µm thick mouse brain coronal slices using a CM3050 S cryostat (Leica, Germany) at −20°C, and thaw-mounted onto indium tin oxide (ITO) coated glass slides (Bruker Daltonics, Germany) [44]. Each treatment group contained four female and four male mice, and four slices from each animal were used for MALDI imaging experiments, half of them containing the dorsal hippocampus (Bregma −1.22 to −1.94 mm) and the other half the ventral hippocampus (Bregma −2.54 to −3.28 mm) [41] (Figure 1A).

After a wash in Carnoy’s solution (6 ethanol: 3 chloroform: 1 acetic acid; all from Thermo Fisher, USA), used to removed most of the lipids from the section [45], the slides were allowed to dry. Then, a solution containing 25 µg/mL of sequencing grade trypsin (Promega, USA) diluted in 20 mM ammonium bicarbonate (Thermo Fisher, USA) was uniformly applied onto the tissue sections using a HTX TM-SprayerTM (HTX Technologies, USA) in 15 layers with a spray flow rate of 10 µL/min at 30°C.

Afterwards, slides were placed in an airtight container in a humid atmosphere and incubated at 37°C overnight, then coated with 7 mg/mL of CHCA matrix (α-cyano-4- hydroxycinnamic acid; Bruker Daltonics, Germany) in a 70% acetonitrile solution (containing 1% trifluoroacetic acid, TFA) using the HTX TM-SprayerTM with the following parameters: temperature 75°C; number of passes 8; flow rate 120 µL/min; velocity 1200 mm/min; track spacing 3 mm; pressure 10 psi (Figure 1B).

### 2.4. MALDI imaging acquisition and analysis

All imaging analysis were performed using a RapifleXTM MALDI TissuetyperTM TOF/TOF mass spectrometer (Bruker Daltonics, Germany) equipped with a Smartbeam™ 3D laser (Figure 1B).

Mass measurements were performed in reflector positive ion mode in the m/z range of 600 to 3500 Da, with a spatial resolution of 50 µm. External calibration was performed using Peptide Calibration Standard Kit II (Bruker Daltonics, Germany). For histological annotation and precise hippocampal delineation, standard Nissl staining was applied to consecutive tissue sections and photographed using an Axio Imager.M2 microscope (Zeiss, Germany) (Figure 1C).

Visualization and statistical analysis were performed with FlexImaging and SCiLS Lab software (2023b version; Bruker Daltonics, Germany). Data from the tissue sections were imported to SCiLS Lab. The processing steps included baseline subtraction (Top-hat filter), normalisation (Total Ion Current algorithm), and spatial denoising (weak). Peaks were aligned to the mean spectrum by centroid matching. Average spectra, representative of the whole measurement regions and ROIs, were generated to display differences in the peptide profiles (Figure 1D). Intensity values for each m/z peak were exported to Excel for further calculations. MetaboAnalyst was used to normalized abundance of each m/z peak by log transformation and to generate the volcano plot.

### 2.5. MS/MS of digested tissue sections

Once each m/z was analysed, the identification of the corresponding protein of the ones that showed significant differences between the groups was performed by spatially targeting and sequencing peptides using traditional tandem mass spectrometry (MS/MS) approaches directly on the tissue. The generated MS/MS spectra were submitted to a MASCOT (Matrix Science, USA) database search engine using BioTools software (Bruker Daltonics, Germany) to match tryptic peptide sequences to their respective intact proteins. The search was performed with a tolerance of 100 ppm and ± 0.3 Da against the *Mus musculus* database.

### 2.6. Study of protein-protein interaction (PPI)

To generate a functional association network of the identified proteins and calculate the protein-protein interaction (PPI) enrichment *p*-value, STRING v11.5 database (https://string-db.org/) was used. A significant PPI value means that the proteins included in the study have more interactions among themselves than what would be expected for a random set of proteins of similar size, drawn from the genome, and indicates that the proteins are partially biologically connected as a group [46]. Statistics were performed using the STRING database statistics package, with a confidence level of 0.4.

### 2.7. Western blot

To prove the reproducibility of our proteomic results, some of the proteins identified by MALDI imaging analysis as significantly altered, as well as a results-related protein of interest, were additionally measured by Western blot.

Hippocampal tissue samples (n = 5-9 per group) were homogenized in ice-cold RIPA lysis buffer (50 mM Tris-HCl pH 7.4; 150 mM NaCl; 0.1% Tx100; 0.5% sodium deoxycholate; 0.1% SDS) with protease and phosphatase inhibitors (all from Roche Diagnostics, Germany). Equal amounts of protein (30 μg) were mixed with Laemmli buffer (Bio-rad, USA), then loaded on a sodium dodecyl-sulfate polyacrylamide gel electrophoresis (SDS-PAGE) and subjected to electrophoresis. Proteins were transferred to nitrocellulose membranes (Bio-rad, USA) by using a transblot apparatus (Bio-Rad, USA). Membranes were blocked with 5% dried skimmed milk powder in Tween-PBS for 1 h. Primary antibodies against each protein of interest were applied at appropriate dilution overnight at 4°C (Table 1). After washing, corresponding secondary antibodies (anti-rabbit or anti-mouse HRP-conjugated, #AP307 and #AP130P respectively; Sigma-Aldrich, US) were added for 1 h at a dilution of 1/5000. Blots were washed, incubated in enhanced chemiluminescence reagent (ECL Prime; Bio-rad, USA), and developed using the G:BOX Chemi XX6 gel documentation system (Syngene, India).

**Table 1.** Characteristics of primary antibodies used to measure protein levels by Western blot.

| Protein | Supplier | Host species | Dilution | Molecular weight |
| --- | --- | --- | --- | --- |
| RCAN1 | Proteintech | Rabbit | 1.2000 | 25-30 kDa |
| GluR5 | Proteintech | Rabbit | 1.2000 | 80 KDa |
| pGSK-3β Ser9 | Proteintech | Mouse | 1.2000 | 48 kDa |
| GSK-3β | Proteintech | Rabbit | 1.2000 | 48 KDa |
| β-actin | Sigma-Aldrich | Mouse | 1.5000 | 42 kDa |

An antibody against *β*-actin (Table 1) was used as loading control. For blot quantification, density of each band was determined using ImageJ software (ImageJ, USA). Values were expressed as a ratio of protein of interest/*β*-actin, or phosphorylated protein/total protein when appropriate, and in percentage of control (vehicle) group (100%).

### 2.8. Data analysis

Data were represented as mean ± SEM and analyzed by two or three-way ANOVA, using treatment and sex or treatment, sex and hippocampal region as factors, respectively, and followed by Tukey’s *post-hoc* analysis. When comparing only two groups, unpaired two-tailed Student t test was used. Statistical significance was set at *p* < 0.05, and a fold change (FC) ≤0.66 (downregulated) or ≥1.5 (upregulated). All analyses were performed using SPSS software v.24 (RRID:SCR_002865; IBM, USA) and GraphPad Prism software v.8.3.1 (RRID:SCR_002798; Dotmatics, USA). Final figures were prepared using CorelDraw X8 Software (RRID:SCR_014235; Corel Corporation, Canada).

## 3. Results

### 3.1. Differentially expressed proteins across the whole hippocampus: early amyloidosis and sexually dimorphic differences in synaptic plasticity-related proteins

MALDI imaging quantification detected 234 peptides, of which 34 showed significant differences (1.5-fold change) between groups due to either treatment, sex, hippocampal area, or an interaction between them (Figure 2A). The volcano plot showed that most of those altered proteins were upregulated (74%) and only a few were downregulated (26%). The specific proteins corresponding to those 34 peptides were identified using MS/MS directly on the tissue (Table 2).

**Figure 2.**
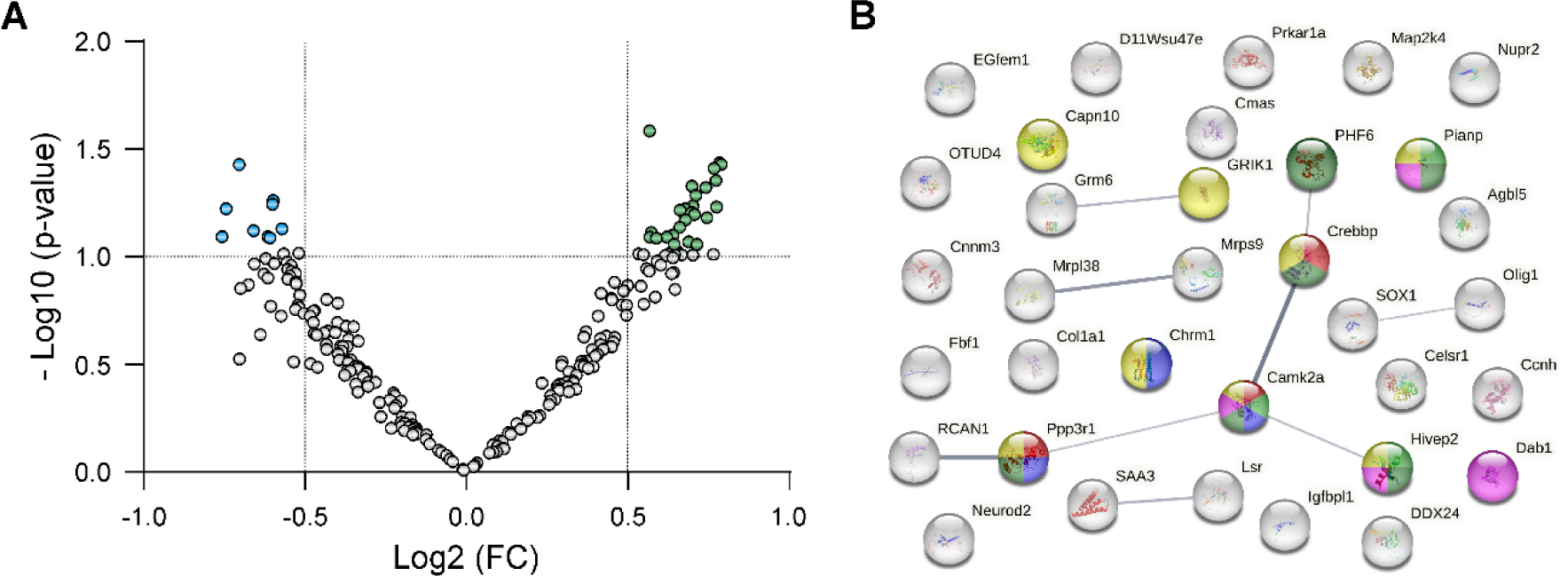
Hippocampal proteins significantly affected by treatment with oA*β*_1-42_, sex or both, and interaction among them. **(A)** Volcano plot showing 9 downregulated (blue points) and 25 upregulated proteins (green points) with a fold change ≥1.5. **(B)** Protein-protein interaction (PPI) of the differentially expressed proteins. A significant PPI (*p* < 0.05), according to STRING database, was found. Each sphere represents a single protein (see details in Table 2). Colored proteins are included in the leading significantly enriched pathway and phenotypes (in order of strength): long term potentiation (red), abnormal response to social novelty (light green), reduced long term depression (blue), abnormal social investigation (dark green), abnormal dentate gyrus morphology (pink) and abnormal synaptic transmission (yellow). Lines represent protein−protein interactions, and line thickness indicates the strength of data supporting the interaction.

**Table 2.**
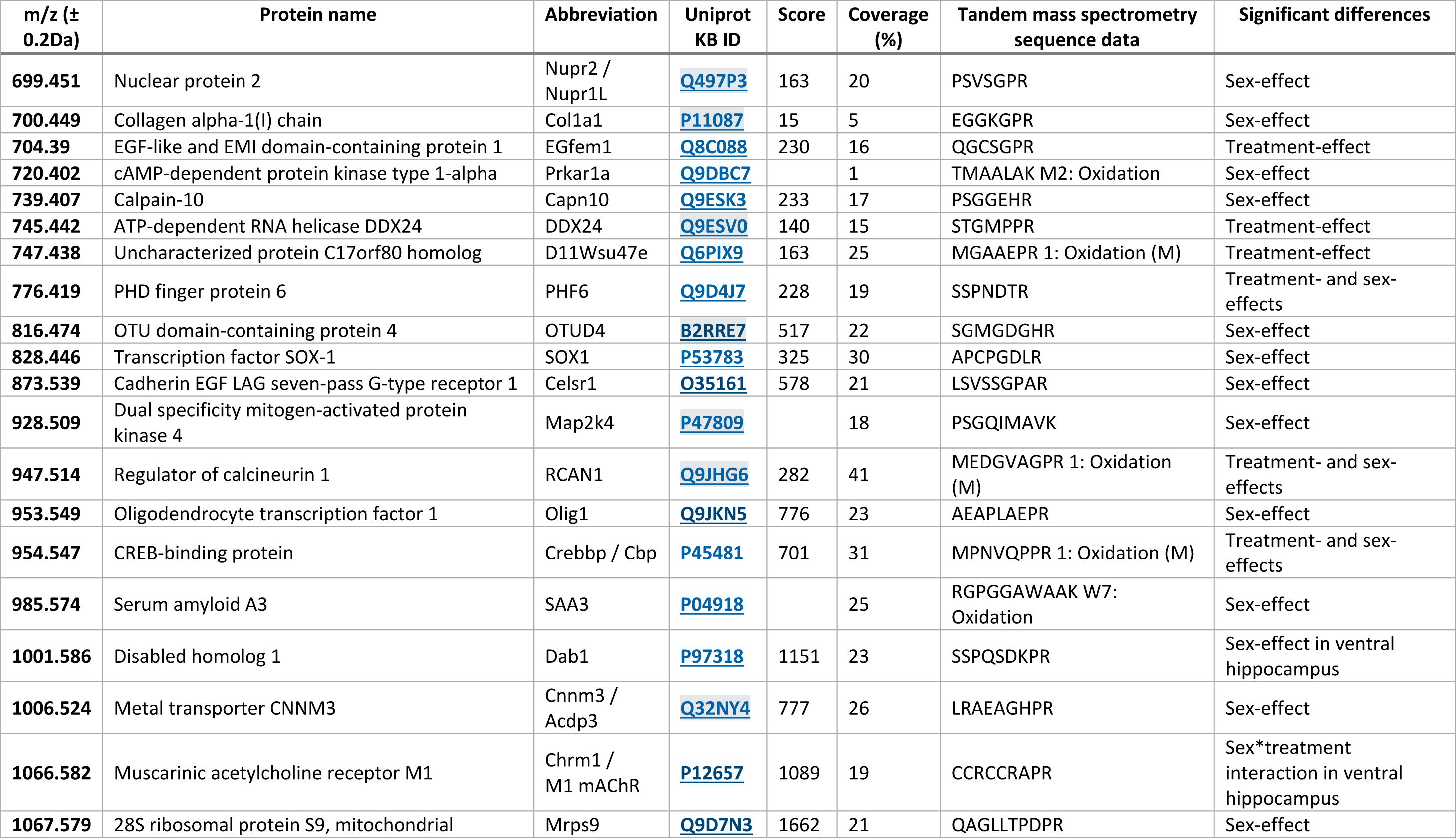
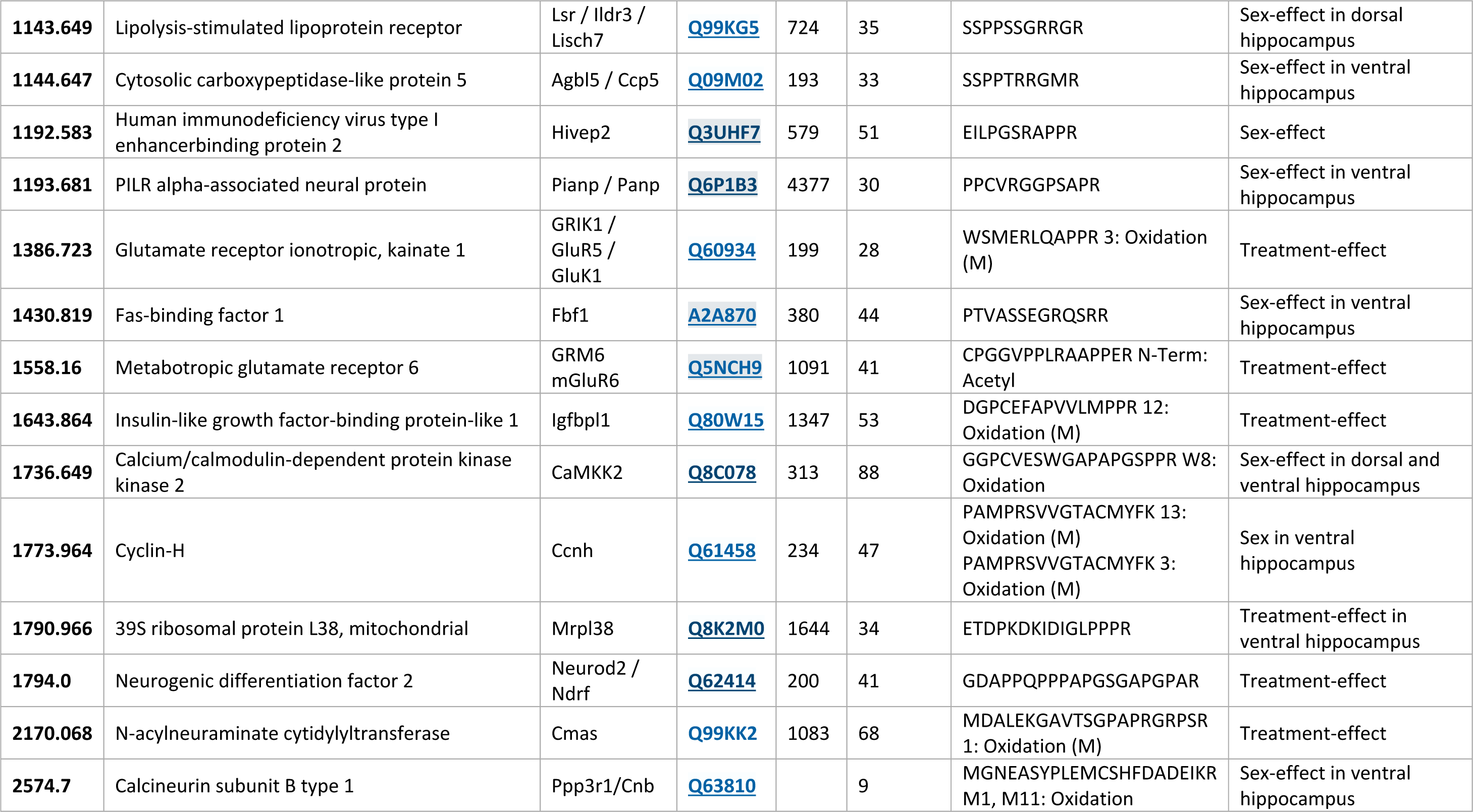
Differentially expressed proteins due to oA*β*_1-42_ treatment and/or sex in the whole, dorsal and ventral hippocampus.

When comparing treated and vehicle groups, differences were found in 13 proteins, while the remaining 21 proteins showed differences when comparing males and females (Table 2).

According to the STRING database, there was a significant interaction (PPI *p*-value = 0.04) among the differentially expressed proteins, being the leading significantly enriched pathway and phenotypes, in order of strength: LTP, abnormal response to social novelty, reduced long term depression (LTD), abnormal social investigation, abnormal dentate gyrus morphology and abnormal synaptic transmission (Figure 2B). Consequently, 14 proteins that after an extensive search of the existing literature were found to be related to synaptic plasticity and/or AD were selected to further study: RCAN1, Crebbp, GluR5, mGluR6, Igfbpl1, Neurod2, SAA3, Hivep2, Lsr, CaMKK2, M1 mAChR, Dab1, Pianp and Ppp3r1.

To map the distribution of the selected 14 proteins across the whole hippocampus, MALDI imaging protein expression data for each protein were analysed (Figure 3). First, the effect of acute hippocampal amyloidosis was studied by analysing changes induced by oA*β*_1-42_ administration. Data showed a downregulation of RCAN1 (treatment effect: F_(2,77)_ = 5.306, *p* = 0.0069) and Crebbp (treatment effect: F_(2,77)_ = 3.777, *p* = 0.0272) due to oA*β*_1-42_ injection. On the other hand, an upregulation of GluR5 (treatment effect: F_(2,79)_ = 4.102, *p* = 0.0202), mGluR6 (treatment effect: F_(2,73)_ = 7.397, *p* = 0.0012), Igfbpl1 (treatment effect: F_(2,78)_ = 6.314, *p* = 0.0029) and Neurod2 (treatment effect: F_(2,80)_ = 6.939, *p* = 0.0017) was observed when comparing treated and control animals.

**Figure 3.**
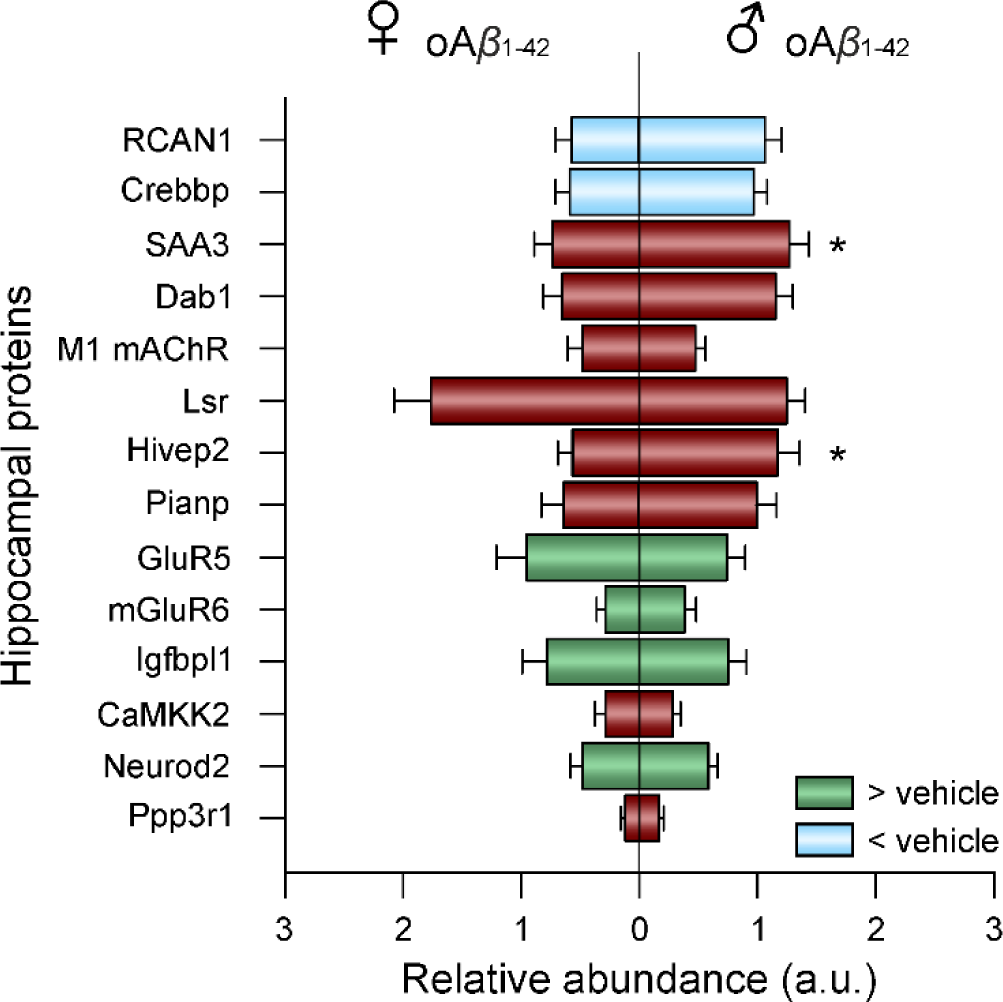
Memory related proteins with significant altered abundance in the whole hippocampus of oA*β*_1-42_ treated female (left) and male (right) mice. Blue indicates downregulation, while green indicates upregulation compared to vehicle of the corresponding sex. Data are expressed as mean ± SEM. * *p* < 0.05 male *vs.* female. A.u., arbitrary units; oA*β*_1-42_, Amyloid-*β_1-42_* oligomers; ♀, female; ♂, male.

Furthermore, regarding potential protein differences due to sexual dimorphism, when comparing oA*β*_1-42_ treated females *vs*. males, differences were found in RCAN1 (sex effect: F_(1,77)_ = 6.956, *p* = 0.0101), SAA3 (sex effect: F_(1,77)_ = 8.501, *p* = 0.0046) and Hivep2 (sex effect: F_(1,51)_ = 4.379, *p* = 0.0414), being all of them higher in males than females. However, none of those proteins showed sex-differences in the vehicle group, which could indicate that those are not physiological sex-differences and it might be a differential effect of oA*β*_1-42_ among males and females.

In order to validate the expressional changes of proteins acquired by the MALDI imaging analyses, Western blot analyses were conducted with two selected altered proteins (RCAN1 and GluR5; Supplementary file 1).

### 3.2. Differentially expressed proteins in the dorsal and ventral hippocampus: early amyloidosis and sexually dimorphic differences in synaptic plasticity- related proteins

To further explore the spatial distribution of the altered proteins taking advantage of this cutting edge technology, data were segmented, separating dorsal and ventral hippocampus, as many studies have shown that there are both molecular and functional differences between the two [38–40].

In the dorsal hippocampus (Figure 4A), two-way ANOVAs revealed treatment effects in the same proteins than showed differences in the whole hippocampus (RCAN1: F_(1,22)_ = 5.281, *p* = 0.0314; Crebbp: F_(1,21)_ = 5.92, *p* = 0.024; GluR5: F_(1,22)_ = 4.211, *p* = 0.05; mGluR6: F_(1,21)_ = 7.348, *p* = 0.0131; Igfbpl1: F_(1,22)_ = 7.535, *p* = 0.0118; Neurod2: F_(1,24)_ = 6.019, *p* = 0.0218), with a downregulation of the first two proteins and an upregulation of the last four proteins in oA*β*_1-42_ mice. Regarding sex effects in this area, differences were found only in Lsr (F_(1,21)_ = 9.575, *p* = 0.0055) and CaMKK2 (F_(1,23)_ = 5.957, *p* = 0.0228) between females and males after oA*β*_1-42_ injection, and none between female and male vehicles.

**Figure 4.**
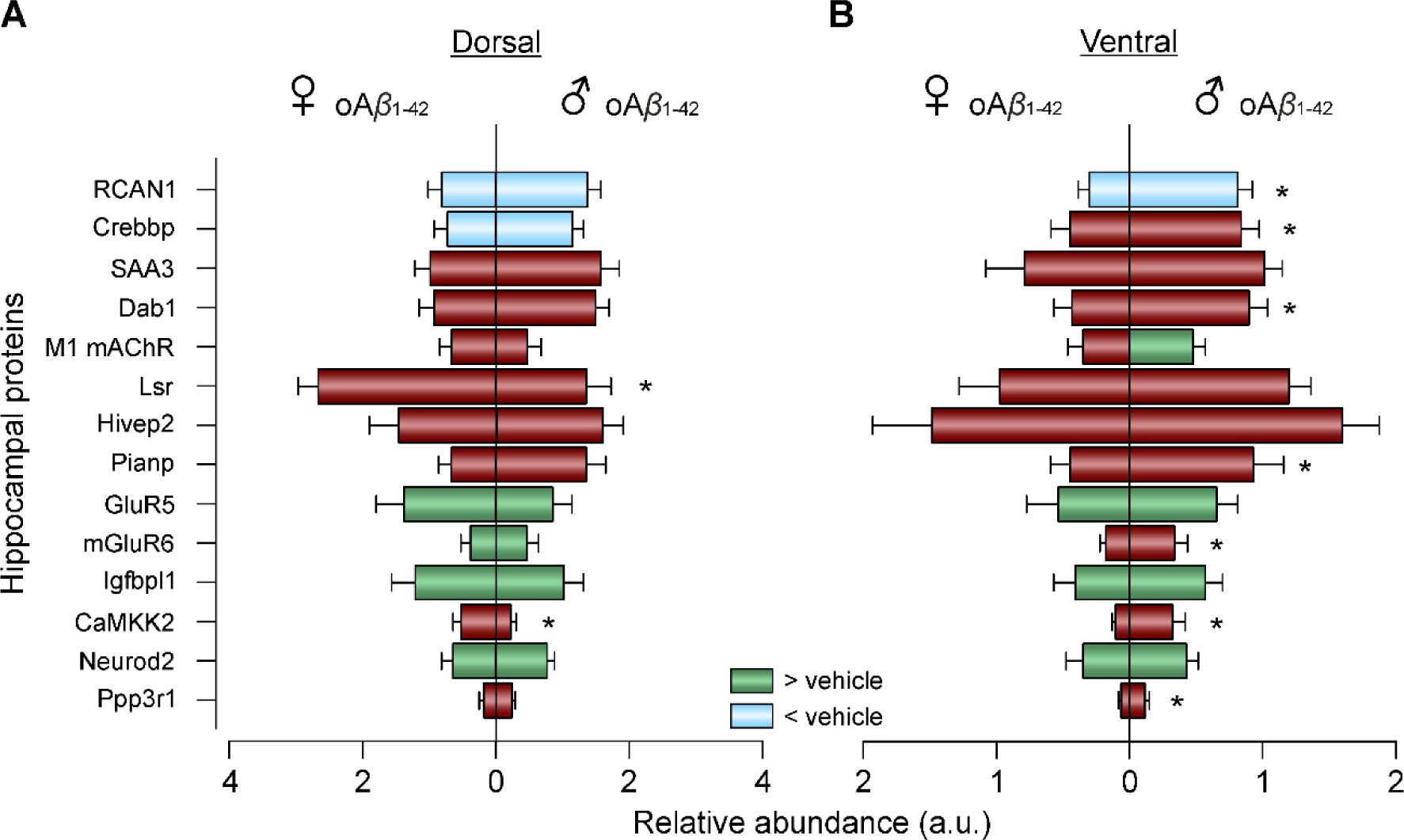
Memory related proteins with significant altered abundance in different hippocampal areas of oA*β*_1-42_-treated female and male mice. Expression of the selected proteins in the dorsal **(A)** and ventral **(B)** hippocampus of oA*β*_1-42_ treated female (left) and male (right) mice. Blue indicates downregulation, while green indicates upregulation compared to vehicle of the corresponding sex. Data are expressed as mean ± SEM. * *p* < 0.05 male *vs.* female. A.u., arbitrary units; oA*β*_1-42_, Amyloid-*β_1-42_* oligomers; ♀, female; ♂, male.

Interestingly, there were less differences due to treatment and more sex effects in the ventral hippocampus (Figure 4B). Specifically, two-way ANOVA revealed a downregulation of RCAN1 (F_(1,24)_ = 6.465, *p* = 0.0179) as well as an upregulation of GluR5 (F_(1,25)_ = 4.521, *p* = 0.0435), Igfbpl1 (F_(1,26)_ = 5.882, *p* = 0.0225) and Neurod2 (F_(1,27)_ = 5.756, *p* = 0.0236) for oA*β*_1-42_-treated mice compared to controls. One protein, M1 mAChR, showed an interaction effect between sex and treatment (F_(1,27)_ = 5.634, *p* = 0.025). Thus, in vehicle animals females showed a higher expression than males (t_13_ = 2.650, *p* = 0.02), however, after oA*β*_1-42_ treatment, that sex-effect was lost and M1 mAChR was upregulated in the male group compared to male controls (t_14_ = 2.395, *p* = 0.031). Moreover, sex differences were found in RCAN1 (F_(1,24)_ = 18.07, *p* = 0.0003), Crebbp (F_(1,27)_ = 7.552, *p* = 0.0106), Dab1 (F_(1,27)_ = 10.65, *p* = 0.003), Pianp (F_(1,27)_ = 5.693, *p* = 0.0243), mGluR6 (F_(1,27)_ = 4.402, *p* = 0.0454), CaMKK2 (F_(1,28)_ = 9.188, *p* = 0.0052) and Ppp3r1 (F_(1,26)_ = 5.496, *p* = 0.027). Once again, they were all specifically between the oA*β*_1-42_ treated males and females, and they were not altered when comparing female and male vehicles.

### 3.3. GSK-3*β* expression and activity alterations due to early amyloidosis treatment

Furthermore, as discussed hereafter, many of the altered protein were found to modulate, either directly or indirectly, glycogen synthase kinase-3*β* (GSK-3*β*), a protein highly involved in the regulation of LTP/LTD induction threshold. Thus, western blot was applied to measure both GSK-3*β* expression and its phosphorylation in Ser9, which causes its inactivation.

Regarding GSK-3*β* expression levels, two-way ANOVA revealed a treatment effect (F_(1,29)_ = 7.281, *p* = 0.0115) and post hoc analysis showed a downregulation of this protein, more pronounced in oA*β*_1-42_ treated males (Figure 5A). Finally, when studying the activity levels of GSK-3*β,* a significant treatment effect was found (F_(1,24)_ = 7.907, *p* = 0.0097), showing a lower phosphorylation of this protein in Ser9 (Figure 5B), especially in female mice. That indicates that even though GSK-3*β* was less expressed in oA*β*_1-42_ treated animals, it was overactivated.

**Figure 5.**
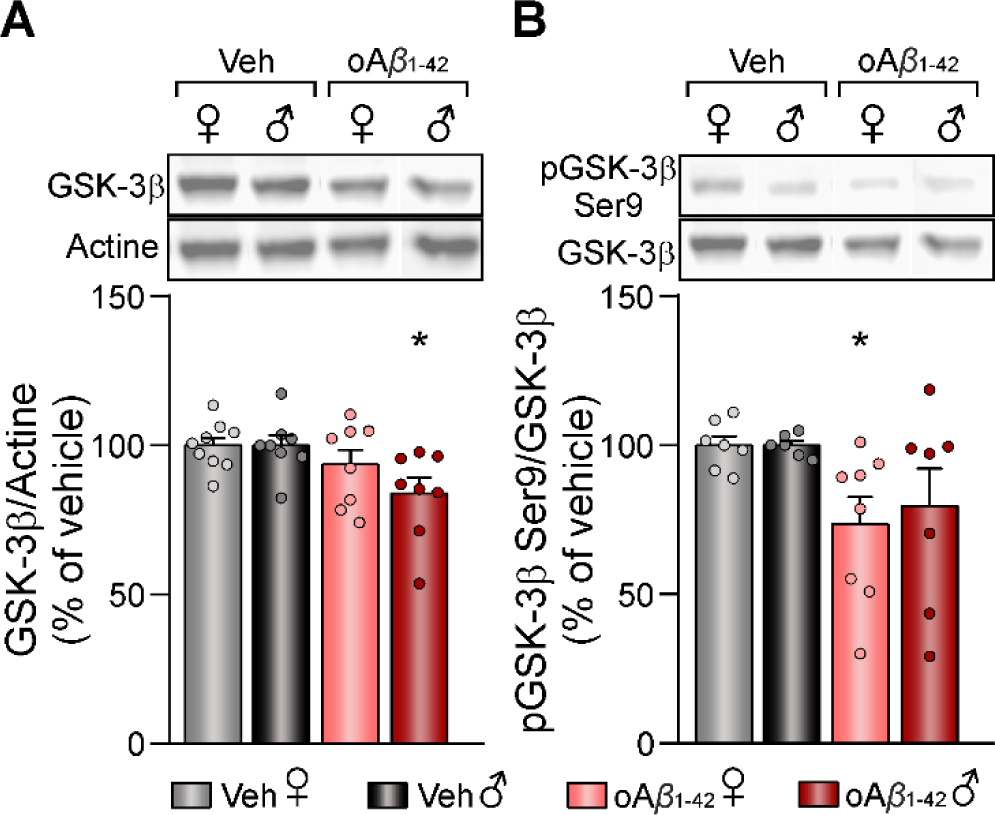
Hippocampal GSK-3*β* expression and activity levels after *icv.* oA*β*_1-42_ treatment. **(A)** Relative expression of GSK-3*β* in vehicle and oA*β*_1-42_ treated mice and representative western blots. Data are expressed as mean ± SEM of the target protein/actine as a loading control, and as percentage (%) of the control of the corresponding sex. **(B)** Relative phosphorylation of GSK-3*β* in Ser9 in vehicle and oA*β*_1-42_ treated mice and representative western blots. Data are expressed as mean ± SEM of the phosphorylated protein/total amount of protein, and as percentage (%) of the control of the corresponding sex. N vehicles: males = 6-8 and females = 7-9; N oA*β*_1-42_: males = 7-8 and females = 8. oA*β*_1-42_, Amyloid-*β_1-42_* oligomers; veh, vehicle; ♀, female; ♂, male. * *p* < 0.05 *vs*. vehicle of the corresponding sex.

## 4. Discussion

MALDI imaging has emerged as an innovative technique, giving the unique opportunity to study protein expression with spatial resolution directly on the tissue. This study describes for the first time the main proteins modified by early amyloidosis in female and male AD mice and provides a mapping of the spatial distribution of those altered proteins in the dorsal and ventral hippocampal areas. Here, we will further discuss the results of a selection of 14 proteins that were found altered due to either oligomeric amyloid treatment or sex, and that are of special interest due to their role in synaptic plasticity and memory.

### Early amyloidosis proteome alterations

Among those 14 proteins, our data showed a downregulation of RCAN1 and Crebbp, and an upregulation of GluR5, mGluR6, Igfbpl1 and Neurod2 in the hippocampus of both male and female mice after a single oA*β*_1-42_ *icv.* injection, a well characterized model of early AD-like amyloidosis [18]. RCAN1 main function is the regulation of the phosphatase calcineurin, which in turn activates or inactivates GSK-3*β* (Figure 6) [47], an effector in the signaling cascade for LTP/LTD induction [48]. Accumulated evidence indicates that RCAN1 expression is chronically elevated in the brains of AD patients, underlying the characteristic neurodegeneration [49, 50]. However, recent data have shown that both loss and gain of RCAN1 in the hippocampus are deleterious, since they promote memory deficits and pathophysiology similar to that observed in AD and Down syndrome patients [51, 52]. Thus, the downregulation of RCAN1 shown in the oA*β*_1-42_ treated mice, could partially account for the imbalance of the LTP/LTD induction threshold previously reported using this amyloidosis murine model [14, 18], without causing neurodegeneration. Moreover, Crebbp is a transcriptional regulator of CREB activity, which also modulates GSK-3*β* expression (Figure 6) [53, 54]. In line with our results, it has been previously reported a low expression of this protein in both AD patients and murine models [53, 55, 56]. Furthermore, restoring Crebbp levels ameliorates learning and memory deficits caused by A*β* [55], pointing to this protein as a potential therapeutic target in AD.

**Figure 6.**
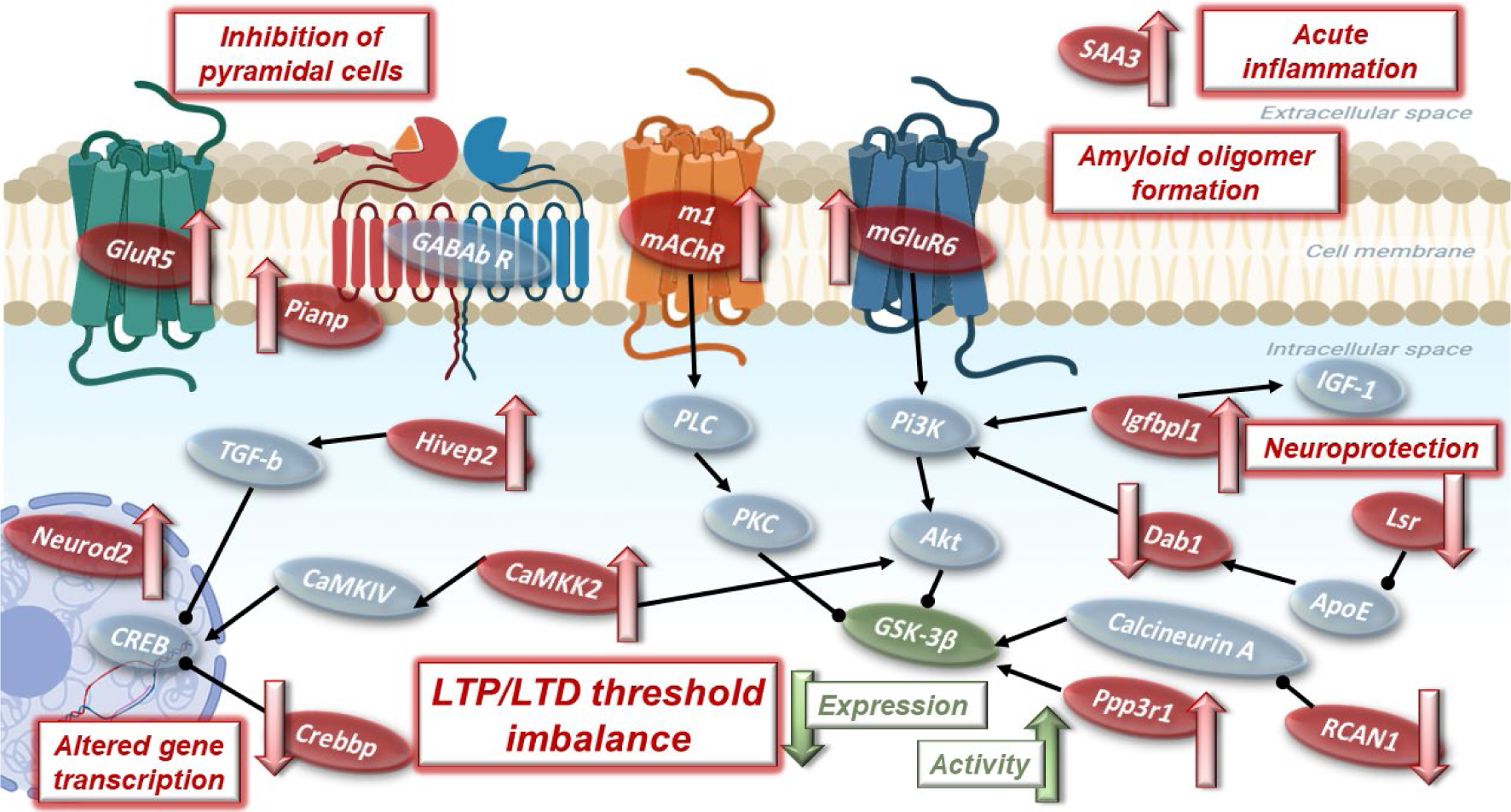
Proposed hippocampal signaling pathway altered by early amyloidosis. Proteins found to be modified by a single oA*β*_1-42_ icv. injection are represented in red, with a red arrow indicating either up- or down-regulation, while proteins on which they exert their effect/modulate are shown in gray and connected with black arrows. Highlighted in green is GSK-3*β* due to its special relevance, as it is modulated by most of the proteins studied, and a decrease in its expression along with an overactivation have been found after oA*β*_1-42_ administration.

Regarding the upregulated proteins, GluR5 is a subtype of kainate receptor that mediate feedforward inhibition of pyramidal cells (Figure 6), raising the threshold for LTP induction [57–59]. Furthermore, this ionotropic receptor is also related to hippocampal *gamma* oscillations [60], which are altered in this murine model of early amyloidosis [16]. Another type of glutamatergic receptor upregulated by *icv.* oA*β*_1-42_ administration is mGluR6. Although this metabotropic glutamate receptor subtype has been typically associated with the retina, recent studies suggest a more widespread expression [61]. In the brain, they are critical for the maintenance of LTP by modulating the production of second messengers via G proteins (Figure 6) [61, 62]. Thus, the higher expression of these two glutamatergic receptors in oA*β*_1-42_ treated mice would also contribute to the LTP deficits observed in this model that subsequently account for robust memory impairments [18]. On the other hand, Igfbpl1 prolongs the half-life of insulin-like growth factor-I (IGF-I) [63], which plays a neuroprotective role (Figure 6) [64, 65]. Thus, the upregulation of Igfbpl1 found in the oA*β*_1-42_ group might be a compensatory way to counteract A*β* neurotoxicity. Moreover, this protein is also involved in the PI3K/AKT signaling, that lead to both LTP and LTD induction [65, 66]. Finally, Neurod2 is a transcription factor involved particularly in AMPA receptor expression (Figure 6) [67]. Enhanced Neurod2 levels, as observed here after oA*β*_1-42_ treatment, have been related to defective glutamatergic synaptic formation and maturation [67]. Therefore, all this results points towards an alteration in GSK-3*β* ability to shift the synaptic plasticity threshold as a potential cause underlying the pathogenesis of early amyloidosis.

Interestingly, most of the protein expression alterations due to oA*β*_1-42_-treatment found when analyzing the whole hippocampus were located in the dorsal hippocampus, which is specifically related to spatial learning and spatial memory [38, 40, 68], but not in the ventral hippocampus, mostly involved in emotional behaviors [38]. This might be explained by the fact that, after *icv*. administration, A*β* diffuses mainly to the dorsal hippocampal formation [14]. The only protein that showed a treatment-effect exclusively in the ventral part of the hippocampus was M1 mAChR, which plays a pivotal role in cognitive and memory processing during sleep (Figure 6) [69]. Moreover, the difference was observed only in male treated mice, in line with previous data from post-mortem brain tissue of AD patients [70]. M1 mAChR is currently being targeted as a promising therapeutic strategy to improve the cognitive decline in AD patients [71]. The alteration of this protein specifically in the ventral hippocampus could indicate a greater involvement of this protein in social memory or anxiety processes associated with this hippocampal area [38], although additional experiments would be needed to further explore this idea.

### Sexual dimorphism differences

Conversely, most of the differences found between male and female mice after oA*β*_1-42_ injection were detected in the ventral hippocampus. In this area, RCAN1 and Crebbp, above discussed, were downregulated specifically in oA*β*_1-42_ treated females. Another protein that was less expressed in the ventral hippocampus of oA*β*_1-42_-treated females compared to males was Dab1, a key regulator of the reelin pathway (Figure 6) that had been previously related to trafficking and processing of APP and apoEr2 [72]. Its phosphorylation leads to the activation of the AKT/GSK-3*β* pathway and the NMDA receptors [73], and both Dab1 expression and phosphorylation are diminished in the cortex and hippocampus of 3xTg-AD mice [74]. These three proteins are downregulated only in the ventral hippocampus of female mice after oA*β*_1-42_ treatment and, since this area is less affected by oA*β* in the AD model used here [14], this sex-differences could be related to the special vulnerability of women to AD [75, 76].

Regarding oA*β*_1-42_-treated males, four proteins were upregulated in the ventral hippocampus: Pianp, mGluR6, CaMKK2 and Ppp3r1. Pianp forms assembles with GABA_B_ receptors and stabilized them in the presynaptic membrane, to regulate synaptic transmission (Figure 6) [77, 78], possibly linking this alteration to hyperexcitability and synaptic plasticity impairments found in early amyloidosis [15, 16]. mGluR6 upregulation, as previously stated, may be related to the deficits in LTP [61]. Another altered protein related to LTP maintenance is CaMKK2. This kinase is necessary for late-LTP, since it is involved in the synthesis of plasticity related proteins and the subsequent memory consolidation (Figure 6) [79–83], and the overactivation of this pathway has been related to synaptic loss in a transgenic AD mice model [84, 85]. Interestingly, CaMKK2 has a male-specific role in hippocampal memory formation [86], thus the overexpression of CaMKK2 in male mice after oA*β*_1-42_ injection might be contributing to LTP and memory deficits specifically in males. Finally, Ppp3r1 is the phosphatase calcineurin B, which activates GSK-3*β* (Figure 6) [47] and a specific polymorphism in the gene encoding this protein is associated with accelerated progression of AD [87–89].

On the other hand, in the dorsal hippocampus only two proteins showed sex-differences. CaMKK2 was the only protein overexpressed in both ventral and dorsal hippocampus, although in the former the alteration was observed in males while in the latter was in the female oA*β*_1-42_ group. Additionally, Lsr expression was lower in oA*β*_1-42_ males than females. The suppression of Lsr has revealed deficits similar to those reported in neurodegenerative diseases, such as impairments in social and visual memory, as well as short-term working memory problems (Figure 6) [90], with a polymorphism in the gene encoding this protein being associated with a higher risk for AD [91].

Thus, it appears that in the ventral hippocampus, where less oA*β*_1-42_ is infused after *icv*. injection, sexual dimorphism is more evident than in the dorsal part of the hippocampus, in which most of the alterations were caused by early amyloidosis regardless the sex.

Finally, considering the whole hippocampus, sex-differences were observed in 2 proteins, both upregulated in males and downregulated in females after oA*β*_1-42_ treatment. The first one, SAA3, is a major acute-phase protein during inflammatory responses, that is implicated in amyloid deposition and colocalized with senile plaques in AD brains (Figure 6) [92, 93]. The alteration in this protein is related to the misfolding and oligomer formation that take place in AD and other amyloidosis diseases [94, 95]. The other altered protein, Hivep2, is a protein related to intellectual disability and specifically implicated in short-term synaptic plasticity (Figure 6) [96, 97]. It is noteworthy that different results are observed when analyzing the whole hippocampus and the dorsal vs. ventral areas of the hippocampus separately, stressing once again the importance of mapping of the spatial proteome distribution in AD.

### GSK-3*β* as a biomarker of synaptic plasticity alterations in early amyloidosis

Interestingly, many of the altered proteins were found to modulate GSK-3*β*, either directly or indirectly, and as revealed by the western blot assay, both its expression and activity levels were indeed altered by a single oA*β*_1-42_ *icv.* injection. This kinase showed a decreased expression level and decreased phosphorylation in the Ser9 residue, which leads to the activation of GSK-3*β.* Its activation has been widely related to a shift in the LTP/LTD induction threshold, tilting it towards an induction of LTD [98]. Hence, this activation of GSK-3*β* in the oA*β*_1-42_ treated mice, caused at least partially by the altered proteins revealed by MALDI imaging analysis, could underlie the transformation of HFS-induced LTP into LTD previously described *ex vivo* [14, 18] and *in vivo* [15, 16] and therefore the memory deficits present in this murine model [18]. This overactivation of GSK-3*β* has been previously reported in AD, and as a result it has become a potential therapeutic target [99] that deserve further investigation.

## 5. Conclusions

A recent work has shown potential biomarkers related to neurodegeneration in the cortex of 5xFAD mice using proteomic MALDI imaging [37]. However, to the best of our knowledge, our work is the first that provides a mapping of the spatial distribution of the hippocampal proteome, which is of special relevance for next-generation *in vivo* modeling of AD [100]. Overall, our results showed alterations in both male and female mice after a single *icv.* oA*β*_1-42_ injection in the expression of several key proteins related to memory formation and the underlying synaptic plasticity LTP/LTD processes [18] (Figure 6), but not neurodegeneration. This endorses the use of this model as a robust way to study the early stages of AD, when neurodegeneration is not yet present. Furthermore, GSK-3*β* arises as a promising biomarker of aberrant plasticity and memory caused by early amyloidosis and as an important molecular therapeutic target.

### Future perspectives

The mapping of the spatial hippocampal proteome facilitated here might provide a specific signature of biomarkers of AD’s early stages. A recent proteomic analysis has demonstrated a regional heterogeneity in the protein expression of the different hippocampal subfields (i.e. CA1, CA2, CA3 and dentate gyrus) in healthy humans [101], and regarding AD, lipids expression seem to change specifically in the CA1 subfield of the hippocampus [102]. Since each subfield is attributed a different function [103–106], the question arises whether similar results to those presented here would be observed regarding synaptic plasticity-related proteins if the hippocampal proteome of each subfield is analyzed independently. Hence, future MALDI imaging studies are needed to map spatial proteomic distribution and changes caused by AD in the different hippocampal subfields and its implications for early diagnosis and treatment.

## List of abbreviations

AD: Alzheimer’s disease
APP: amyloid precursor protein
A*β*: amyloid-*β*
oA*β*_1-42_: amyloid-*β* oligomers
E/I: excitatory/inhibitory
GSK-3*β*: glycogen synthase kinase-3*β*
*Icv*: Intracerebroventricularly
IGF-I: insulin-like growth factor-I
LTD: long-term depression
LTP: long-term potentiation
MALDI: matrix-assisted laser desorption/ionization
MS/MS: tandem mass spectrometry
PBS: Phosphate-buffered saline
PPI: protein-protein interaction

## Declarations

### Consent for publication

Not applicable.

### Availability of data and materials

The datasets used and/or analyzed during the current study are available from the corresponding authors on reasonable request.

## Authors’ contributions

LJD and JDNL were responsible for the initial conceptualization; RJH and SD performed the surgeries, drug administration and sacrifice; AC performed the proteomic assay; AC analyzed the data; AC was responsible for writing the original draft; AC, JDNL and LJD were responsible for visualization and did the writing – review and editing. LJD and JDNL were responsible for funding acquisition, supervision, and project administration. All authors read and approved the final manuscript.

## Acknowledgements

We acknowledge Dr. Pilar Alberdi for her excellent technical help with the MALDI mass spectrometer.

## References

1. Gauthier S, Rosa-Neto P, Morais JA, Webster C: World Alzheimer Report 2021: Journey through the diagnosis of dementia. In.; 2021.

2. Selkoe DJ, Hardy J: The amyloid hypothesis of Alzheimer’s disease at 25 years. EMBO Mol Med 2016, 8(6):595–608.

3. Jeremic D, Jiménez-Díaz L, Navarro-López JD: Past, present and future of therapeutic strategies against amyloid-β peptides in Alzheimer’s disease: a systematic review. Ageing Res Rev 2021, 72:101496.

4. Vyas Y, Montgomery JM, Cheyne JE: Hippocampal Deficits in Amyloid-β-Related Rodent Models of Alzheimer’s Disease. Front Neurosci 2020, 14:266.

5. Palop JJ, Mucke L: Amyloid-beta-induced neuronal dysfunction in Alzheimer’s disease: from synapses toward neural networks. Nat Neurosci 2010, 13(7):812–818.

6. Lührs T, Ritter C, Adrian M, Riek-Loher D, Bohrmann B, Döbeli H, Schubert D, Riek R: 3D structure of Alzheimer’s amyloid-β(1–42) fibrils. Proc Natl Acad Sci U S A 2005, 102(48):17342–17347.

7. Sanchez-Varo R, Mejias-Ortega M, Fernandez-Valenzuela JJ, Nuñez-Diaz C, Caceres-Palomo L, Vegas-Gomez L, Sanchez-Mejias E, Trujillo-Estrada L, Garcia-Leon JA, Moreno-Gonzalez I et al: Transgenic Mouse Models of Alzheimer’s Disease: An Integrative Analysis. Int J Mol Sci 2022, 23(10).

8. Braak H, Thal DR, Ghebremedhin E, Del Tredici K: Stages of the pathologic process in Alzheimer disease: age categories from 1 to 100 years. J Neuropathol Exp Neurol 2011, 70(11):960–969.

9. Michno W, Wehrli P, Meier SR, Sehlin D, Syvänen S, Zetterberg H, Blennow K, Hanrieder J: Chemical imaging of evolving amyloid plaque pathology and associated Aβ peptide aggregation in a transgenic mouse model of Alzheimer’s disease. J Neurochem 2020, 152(5):602–616.

10. Tamagno E, Guglielmotto M, Monteleone D, Manassero G, Vasciaveo V, Tabaton M: The Unexpected Role of Aβ1-42 Monomers in the Pathogenesis of Alzheimer’s Disease. J Alzheimers Dis 2018, 62(3):1241–1245.

11. Mucke L, Selkoe DJ: Neurotoxicity of amyloid β-protein: synaptic and network dysfunction. Cold Spring Harb Perspect Med 2012, 2(7):a006338.

12. Kent SA, Spires-Jones TL, Durrant CS: The physiological roles of tau and Aβ: implications for Alzheimer’s disease pathology and therapeutics. Acta Neuropathol 2020, 140(4):417–447.

13. Li S, Selkoe DJ: A mechanistic hypothesis for the impairment of synaptic plasticity by soluble Abeta oligomers from Alzheimer’s brain. J Neurochem 2020, 154(6):583–597.

14. Sanchez-Rodriguez I, Djebari S, Temprano-Carazo S, Vega-Avelaira D, Jimenez-Herrera R, Iborra-Lazaro G, Yajeya J, Jimenez-Diaz L, Navarro-Lopez JD: Hippocampal long-term synaptic depression and memory deficits induced in early amyloidopathy are prevented by enhancing G-protein-gated inwardly rectifying potassium channel activity. J Neurochem 2020, 153(3):362–376.

15. Sanchez-Rodriguez I, Gruart A, Delgado-Garcia JM, Jimenez-Diaz L, Navarro-Lopez JD: Role of GirK Channels in Long-Term Potentiation of Synaptic Inhibition in an In Vivo Mouse Model of Early Amyloid-beta Pathology. Int J Mol Sci 2019, 20(5).

16. Sánchez-Rodríguez I, Temprano-Carazo S, Nájera A, Djebari S, Yajeya J, Gruart A, Delgado-García JM, Jiménez-Díaz L, Navarro-López JD: Activation of G-protein-gated inwardly rectifying potassium (Kir3/GirK) channels rescues hippocampal functions in a mouse model of early amyloid-β pathology. Sci Rep 2017, 7(1):14658.

17. Morroni F, Sita G, Tarozzi A, Rimondini R, Hrelia P: Early effects of Aβ1-42 oligomers injection in mice: Involvement of PI3K/Akt/GSK3 and MAPK/ERK1/2 pathways. Behav Brain Res 2016, 314:106–115.

18. Jiménez-Herrera R, Contreras A, Djebari S, Mulero-Franco J, Iborra-Lázaro G, Jeremic D, Navarro-López J, Jiménez-Díaz L: Systematic characterization of a non-transgenic Aβ(1-42) amyloidosis model: synaptic plasticity and memory deficits in female and male mice. Biol Sex Differ 2023, 14(1):59.

19. Takousis P, Sadlon A, Schulz J, Wohlers I, Dobricic V, Middleton L, Lill CM, Perneczky R, Bertram L: Differential expression of microRNAs in Alzheimer’s disease brain, blood, and cerebrospinal fluid. Alzheimers Dement 2019, 15(11):1468–1477.

20. Bai B, Vanderwall D, Li Y, Wang X, Poudel S, Wang H, Dey KK, Chen PC, Yang K, Peng J: Proteomic landscape of Alzheimer’s Disease: novel insights into pathogenesis and biomarker discovery. Mol Neurodegener 2021, 16(1):55.

21. Zakharova NV, Bugrova AE, Indeykina MI, Fedorova YB, Kolykhalov IV, Gavrilova SI, Nikolaev EN, Kononikhin AS: Proteomic Markers and Early Prediction of Alzheimer’s Disease. Biochemistry (Mosc) 2022, 87(8):762–776.

22. Horgusluoglu E, Neff R, Song WM, Wang M, Wang Q, Arnold M, Krumsiek J, Galindo-Prieto B, Ming C, Nho K et al: Integrative metabolomics-genomics approach reveals key metabolic pathways and regulators of Alzheimer’s disease. Alzheimers Dement 2022, 18(6):1260–1278.

23. Jiang Y, Zhou X, Ip FC, Chan P, Chen Y, Lai NCH, Cheung K, Lo RMN, Tong EPS, Wong BWY et al: Large-scale plasma proteomic profiling identifies a high-performance biomarker panel for Alzheimer’s disease screening and staging. Alzheimers Dement 2022, 18(1):88–102.

24. Oláh Z, Kálmán J, Tóth ME, Zvara Á, Sántha M, Ivitz E, Janka Z, Pákáski M: Proteomic analysis of cerebrospinal fluid in Alzheimer’s disease: wanted dead or alive. J Alzheimers Dis 2015, 44(4):1303–1312.

25. Manavalan A, Mishra M, Feng L, Sze SK, Akatsu H, Heese K: Brain site-specific proteome changes in aging-related dementia. Exp Mol Med 2013, 45(9):e39.

26. Andreev VP, Petyuk VA, Brewer HM, Karpievitch YV, Xie F, Clarke J, Camp D, Smith RD, Lieberman AP, Albin RL et al: Label-free quantitative LC-MS proteomics of Alzheimer’s disease and normally aged human brains. J Proteome Res 2012, 11(6):3053–3067.

27. Yang H, Wittnam JL, Zubarev RA, Bayer TA: Shotgun brain proteomics reveals early molecular signature in presymptomatic mouse model of Alzheimer’s disease. J Alzheimers Dis 2013, 37(2):297–308.

28. Gurel B, Cansev M, Koc C, Ocalan B, Cakir A, Aydin S, Kahveci N, Ulus IH, Sahin B, Basar MK et al: Proteomics Analysis of CA1 Region of the Hippocampus in Pre-, Progression and Pathological Stages in a Mouse Model of the Alzheimer’s Disease. Curr Alzheimer Res 2019, 16(7):613–621.

29. Ismail R, Parbo P, Madsen LS, Hansen AK, Hansen KV, Schaldemose JL, Kjeldsen PL, Stokholm MG, Gottrup H, Eskildsen SF et al: The relationships between neuroinflammation, beta-amyloid and tau deposition in Alzheimer’s disease: a longitudinal PET study. J Neuroinflammation 2020, 17(1):151.

30. Su L, Surendranathan A, Huang Y, Bevan-Jones WR, Passamonti L, Hong YT, Arnold R, Rodríguez PV, Wang Y, Mak E et al: Relationship between tau, neuroinflammation and atrophy in Alzheimer’s disease: The NIMROD study. Information Fusion 2021, 67:116–124.

31. Ikegawa M, Kakuda N, Miyasaka T, Toyama Y, Nirasawa T, Minta K, Hanrieder J: Mass Spectrometry Imaging in Alzheimer’s Disease. Brain Connect 2023.

32. Kim C-M, Diez I, Bueichekú E, Ahn S, Montal V, Sepulcre J: Spatiotemporal Correlation between Amyloid and Tau Accumulations Underlies Cognitive Changes in Aging. The Journal of Neuroscience 2024, 44(7):e0488232023.

33. Chen Y, Hu D, Zhao L, Tang W, Li B: Unraveling metabolic alterations in transgenic mouse model of Alzheimer’s disease using MALDI MS imaging with 4- aminocinnoline-3-carboxamide matrix. Analytica Chimica Acta 2022, 1192:339337.

34. Schubert KO, Weiland F, Baune BT, Hoffmann P: The use of MALDI-MSI in the investigation of psychiatric and neurodegenerative disorders: A review. Proteomics 2016, 16(11-12):1747–1758.

35. Esteve C, Jones EA, Kell DB, Boutin H, McDonnell LA: Mass spectrometry imaging shows major derangements in neurogranin and in purine metabolism in the triple- knockout 3×Tg Alzheimer mouse model. Biochim Biophys Acta Proteins Proteom 2017, 1865(7):747–754.

36. Hawkinson TR, Clarke HA, Young LEA, Conroy LR, Markussen KH, Kerch KM, Johnson LA, Nelson PT, Wang C, Allison DB et al: In situ spatial glycomic imaging of mouse and human Alzheimer’s disease brains. Alzheimers Dement 2022, 18(10):1721–1735.

37. Uras I, Karayel-Basar M, Sahin B, Baykal AT: Detection of early proteomic alterations in 5xFAD Alzheimer’s disease neonatal mouse model via MALDI-MSI. Alzheimers Dement 2023.

38. Fanselow MS, Dong HW: Are the dorsal and ventral hippocampus functionally distinct structures? Neuron 2010, 65(1):7–19.

39. Floriou-Servou A, von Ziegler L, Stalder L, Sturman O, Privitera M, Rassi A, Cremonesi A, Thöny B, Bohacek J: Distinct Proteomic, Transcriptomic, and Epigenetic Stress Responses in Dorsal and Ventral Hippocampus. Biol Psychiatry 2018, 84(7):531–541.

40. Lee AR, Kim JH, Cho E, Kim M, Park M: Dorsal and Ventral Hippocampus Differentiate in Functional Pathways and Differentially Associate with Neurological Disease- Related Genes during Postnatal Development. Front Mol Neurosci 2017, 10:331.

41. Paxinos G, Franklin KBJ: The Mouse Brain in Stereotaxic Coordinates: Elsevier Academic Press; 2004.

42. Jimenez-Herrera R, Contreras A, Djebari S, Mulero-Franco J, Iborra-Lazaro G, Jeremic D, Navarro-Lopez J, Jimenez-Diaz L: Systematic characterization of a non-transgenic Abeta(1-42) amyloidosis model: synaptic plasticity and memory deficits in female and male mice. Biol Sex Differ 2023, 14(1):59.

43. Pavliukeviciene B, Zentelyte A, Jankunec M, Valiuliene G, Talaikis M, Navakauskiene R, Niaura G, Valincius G: Amyloid β oligomers inhibit growth of human cancer cells. PLoS One 2019, 14(9):e0221563.

44. Høiem TS, Andersen MK, Martin-Lorenzo M, Longuespée R, Claes BSR, Nordborg A, Dewez F, Balluff B, Giampà M, Sharma A et al: An optimized MALDI MSI protocol for spatial detection of tryptic peptides in fresh frozen prostate tissue. Proteomics 2022, 22(10):e2100223.

45. Deutskens F, Yang J, Caprioli RM: High spatial resolution imaging mass spectrometry and classical histology on a single tissue section. J Mass Spectrom 2011, 46(6):568–571.

46. Szklarczyk D, Gable AL, Nastou KC, Lyon D, Kirsch R, Pyysalo S, Doncheva NT, Legeay M, Fang T, Bork P et al: The STRING database in 2021: customizable protein-protein networks, and functional characterization of user-uploaded gene/measurement sets. Nucleic Acids Res 2021, 49(D1):D605–d612.

47. Kim Y, Lee YI, Seo M, Kim SY, Lee JE, Youn HD, Kim YS, Juhnn YS: Calcineurin dephosphorylates glycogen synthase kinase-3 beta at serine-9 in neuroblast-derived cells. J Neurochem 2009, 111(2):344–354.

48. Dudilot A, Trillaud-Doppia E, Boehm J: RCAN1 Regulates Bidirectional Synaptic Plasticity. Curr Biol 2020, 30(7):1167–1176.e1162.

49. Ermak G, Davies KJA: Chronic high levels of the RCAN1-1 protein may promote neurodegeneration and Alzheimer disease. Free Radic Biol Med 2013, 62:47–51.

50. Wu Y, Ly PT, Song W: Aberrant expression of RCAN1 in Alzheimer’s pathogenesis: a new molecular mechanism and a novel drug target. Mol Neurobiol 2014, 50(3):1085–1097.

51. Lee SK, Ahnn J: Regulator of Calcineurin (RCAN): Beyond Down Syndrome Critical Region. Mol Cells 2020, 43(8):671–685.

52. Wong H, Buck JM, Borski C, Pafford JT, Keller BN, Milstead RA, Hanson JL, Stitzel JA, Hoeffer CA: RCAN1 knockout and overexpression recapitulate an ensemble of rest-activity and circadian disruptions characteristic of Down syndrome, Alzheimer’s disease, and normative aging. J Neurodev Disord 2022, 14(1):33.

53. Rouaux C, Jokic N, Mbebi C, Boutillier S, Loeffler JP, Boutillier AL: Critical loss of CBP/p300 histone acetylase activity by caspase-6 during neurodegeneration. Embo j 2003, 22(24):6537–6549.

54. Grimes CA, Jope RS: CREB DNA binding activity is inhibited by glycogen synthase kinase-3 beta and facilitated by lithium. J Neurochem 2001, 78(6):1219–1232.

55. Caccamo A, Maldonado MA, Bokov AF, Majumder S, Oddo S: CBP gene transfer increases BDNF levels and ameliorates learning and memory deficits in a mouse model of Alzheimer’s disease. Proc Natl Acad Sci U S A 2010, 107(52):22687–22692.

56. Pláteník J, Fišar Z, Buchal R, Jirák R, Kitzlerová E, Zvěřová M, Raboch J: GSK3β, CREB, and BDNF in peripheral blood of patients with Alzheimer’s disease and depression. Prog Neuropsychopharmacol Biol Psychiatry 2014, 50:83–93.

57. Lerma J, Marques JM: Kainate receptors in health and disease. Neuron 2013, 80(2):292–311.

58. Dhingra S, Yadav J, Kumar J: Structure, Function, and Regulation of the Kainate Receptor. Subcell Biochem 2022, 99:317–350.

59. Nisticò R, Dargan S, Fitzjohn SM, Lodge D, Jane DE, Collingridge GL, Bortolotto ZA: GLUK1 receptor antagonists and hippocampal mossy fiber function. Int Rev Neurobiol 2009, 85:13–27.

60. Fisahn A, Contractor A, Traub RD, Buhl EH, Heinemann SF, McBain CJ: Distinct roles for the kainate receptor subunits GluR5 and GluR6 in kainate-induced hippocampal gamma oscillations. J Neurosci 2004, 24(43):9658–9668.

61. Palazzo E, Boccella S, Marabese I, Pierretti G, Guida F, Maione S: The Cold Case of Metabotropic Glutamate Receptor 6: Unjust Detention in the Retina? Curr Neuropharmacol 2020, 18(2):120–125.

62. Nakanishi S, Nakajima Y, Masu M, Ueda Y, Nakahara K, Watanabe D, Yamaguchi S, Kawabata S, Okada M: Glutamate receptors: brain function and signal transduction. Brain Res Brain Res Rev 1998, 26(2-3):230–235.

63. Allard JB, Duan C: IGF-Binding Proteins: Why Do They Exist and Why Are There So Many? Front Endocrinol (Lausanne) 2018, 9:117.

64. Akterin S, Cowburn RF, Miranda-Vizuete A, Jiménez A, Bogdanovic N, Winblad B, Cedazo-Minguez A: Involvement of glutaredoxin-1 and thioredoxin-1 in beta-amyloid toxicity and Alzheimer’s disease. Cell Death Differ 2006, 13(9):1454–1465.

65. Wei W, Wang X, Kusiak JW: Signaling events in amyloid beta-peptide-induced neuronal death and insulin-like growth factor I protection. J Biol Chem 2002, 277(20):17649–17656.

66. Guo C, Cho KS, Li Y, Tchedre K, Antolik C, Ma J, Chew J, Utheim TP, Huang XA, Yu H et al: IGFBPL1 Regulates Axon Growth through IGF-1-mediated Signaling Cascades. Sci Rep 2018, 8(1):2054.

67. Safari MS, Obexer D, Baier-Bitterlich G, Zur Nedden S: PKN1 Is a Novel Regulator of Hippocampal GluA1 Levels. Front Synaptic Neurosci 2021, 13:640495.

68. Moser MB, Moser EI, Forrest E, Andersen P, Morris RG: Spatial learning with a minislab in the dorsal hippocampus. Proc Natl Acad Sci U S A 1995, 92(21):9697–9701.

69. Dwomoh L, Tejeda GS, Tobin AB: Targeting the M1 muscarinic acetylcholine receptor in Alzheimer’s disease. Neuronal Signal 2022, 6(1):Ns20210004.

70. Sanfilippo C, Giuliano L, Castrogiovanni P, Imbesi R, Ulivieri M, Fazio F, Blennow K, Zetterberg H, Di Rosa M: Sex, Age, and Regional Differences in CHRM1 and CHRM3 Genes Expression Levels in the Human Brain Biopsies: Potential Targets for Alzheimer’s Disease-related Sleep Disturbances. Curr Neuropharmacol 2023, 21(3):740–760.

71. Scarpa M, Hesse S, Bradley SJ: M1 muscarinic acetylcholine receptors: A therapeutic strategy for symptomatic and disease-modifying effects in Alzheimer’s disease? Adv Pharmacol 2020, 88:277–310.

72. Hoe HS, Tran TS, Matsuoka Y, Howell BW, Rebeck GW: DAB1 and Reelin effects on amyloid precursor protein and ApoE receptor 2 trafficking and processing. J Biol Chem 2006, 281(46):35176–35185.

73. Bracher-Smith M, Leonenko G, Baker E, Crawford K, Graham AC, Salih DA, Howell BW, Hardy J, Escott-Price V: Whole genome analysis in APOE4 homozygotes identifies the DAB1-RELN pathway in Alzheimer’s disease pathogenesis. Neurobiol Aging 2022, 119:67–76.

74. Mota SI, Ferreira IL, Valero J, Ferreiro E, Carvalho AL, Oliveira CR, Rego AC: Impaired Src signaling and post-synaptic actin polymerization in Alzheimer’s disease mice hippocampus--linking NMDA receptors and the reelin pathway. Exp Neurol 2014, 261:698–709.

75. Viña J, Lloret A: Why women have more Alzheimer’s disease than men: gender and mitochondrial toxicity of amyloid-beta peptide. J Alzheimers Dis 2010, 20 Suppl 2:S527–533.

76. 2022 Alzheimer’s disease facts and figures. Alzheimers Dement 2022, 18(4):700-789.

77. Winkler M, Biswas S, Berger SM, Küchler M, Preisendörfer L, Choo M, Früh S, Rem PD, Enkel T, Arnold B et al: Pianp deficiency links GABA(B) receptor signaling and hippocampal and cerebellar neuronal cell composition to autism-like behavior. Mol Psychiatry 2020, 25(11):2979–2993.

78. Dinamarca MC, Raveh A, Schneider A, Fritzius T, Früh S, Rem PD, Stawarski M, Lalanne T, Turecek R, Choo M et al: Complex formation of APP with GABA(B) receptors links axonal trafficking to amyloidogenic processing. Nat Commun 2019, 10(1):1331.

79. Redondo RL, Okuno H, Spooner PA, Frenguelli BG, Bito H, Morris RG: Synaptic tagging and capture: differential role of distinct calcium/calmodulin kinases in protein synthesis-dependent long-term potentiation. J Neurosci 2010, 30(14):4981–4989.

80. Redondo RL, Morris RG: Making memories last: the synaptic tagging and capture hypothesis. Nat Rev Neurosci 2011, 12(1):17–30.

81. Moncada D, Ballarini F, Viola H: Behavioral Tagging: A Translation of the Synaptic Tagging and Capture Hypothesis. Neural Plast 2015, 2015:650780.

82. Okuda K, Højgaard K, Privitera L, Bayraktar G, Takeuchi T: Initial memory consolidation and the synaptic tagging and capture hypothesis. Eur J Neurosci 2021, 54(8):6826–6849.

83. Nonaka M, Fujii H, Kim R, Kawashima T, Okuno H, Bito H: Untangling the two-way signalling route from synapses to the nucleus, and from the nucleus back to the synapses. Philos Trans R Soc Lond B Biol Sci 2014, 369(1633):20130150.

84. Lee A, Kondapalli C, Virga DM, Lewis TL, Jr., Koo SY, Ashok A, Mairet-Coello G, Herzig S, Foretz M, Viollet B et al: Aβ42 oligomers trigger synaptic loss through CAMKK2- AMPK-dependent effectors coordinating mitochondrial fission and mitophagy. Nat Commun 2022, 13(1):4444.

85. Wang L, Yu C, Tao Y, Yang X, Jiang Q, Yu H, Zhang J: Transcriptome analysis reveals potential marker genes for diagnosis of Alzheimer’s disease and vascular dementia. Front Genet 2022, 13:1038585.

86. Mizuno K, Antunes-Martins A, Ris L, Peters M, Godaux E, Giese KP: Calcium/calmodulin kinase kinase beta has a male-specific role in memory formation. Neuroscience 2007, 145(2):393–402.

87. Peterson D, Munger C, Crowley J, Corcoran C, Cruchaga C, Goate AM, Norton MC, Green RC, Munger RG, Breitner JC et al: Variants in PPP3R1 and MAPT are associated with more rapid functional decline in Alzheimer’s disease: the Cache County Dementia Progression Study. Alzheimers Dement 2014, 10(3):366–371.

88. Robinson RA, Lange MB, Sultana R, Galvan V, Fombonne J, Gorostiza O, Zhang J, Warrier G, Cai J, Pierce WM et al: Differential expression and redox proteomics analyses of an Alzheimer disease transgenic mouse model: effects of the amyloid-β peptide of amyloid precursor protein. Neuroscience 2011, 177:207–222.

89. Zhou Z, Bai J, Zhong S, Zhang R, Kang K, Zhang X, Xu Y, Zhao C, Zhao M: Integrative genomic analysis of PPP3R1 in Alzheimer’s disease: a potential biomarker for predictive, preventive, and personalized medical approach. Epma j 2021, 12(4):647–658.

90. El Hajj A, Herzine A, Calcagno G, Désor F, Djelti F, Bombail V, Denis I, Oster T, Malaplate C, Vigier M et al: Targeted Suppression of Lipoprotein Receptor LSR in Astrocytes Leads to Olfactory and Memory Deficits in Mice. Int J Mol Sci 2022, 23(4).

91. Petrelis AM, Stathopoulou MG, Kafyra M, Murray H, Masson C, Lamont J, Fitzgerald P, Dedoussis G, Yen FT, Visvikis-Siest S: VEGF-A-related genetic variants protect against Alzheimer’s disease. Aging (Albany NY) 2022, 14(6):2524–2536.

92. Lin A, Liu J, Gong P, Chen Y, Zhang H, Zhang Y, Yu Y: Serum amyloid A inhibits astrocyte migration via activating p38 MAPK. J Neuroinflammation 2020, 17(1):254.

93. Liu J, Wang D, Li SQ, Yu Y, Ye RD: Suppression of LPS-induced tau hyperphosphorylation by serum amyloid A. J Neuroinflammation 2016, 13:28.

94. Jayaraman S, Gantz DL, Haupt C, Gursky O: Serum amyloid A forms stable oligomers that disrupt vesicles at lysosomal pH and contribute to the pathogenesis of reactive amyloidosis. Proc Natl Acad Sci U S A 2017, 114(32):E6507–e6515.

95. Patke S, Srinivasan S, Maheshwari R, Srivastava SK, Aguilera JJ, Colón W, Kane RS: Characterization of the oligomerization and aggregation of human Serum Amyloid A. PLoS One 2013, 8(6):e64974.

96. Kobayashi K, Takagi T, Ishii S, Suzuki H, Miyakawa T: Attenuated bidirectional short- term synaptic plasticity in the dentate gyrus of Schnurri-2 knockout mice, a model of schizophrenia. Mol Brain 2018, 11(1):56.

97. Srivastava S, Engels H, Schanze I, Cremer K, Wieland T, Menzel M, Schubach M, Biskup S, Kreiß M, Endele S et al: Loss-of-function variants in HIVEP2 are a cause of intellectual disability. Eur J Hum Genet 2016, 24(4):556–561.

98. Jaworski T, Banach-Kasper E, Gralec K: GSK-3β at the Intersection of Neuronal Plasticity and Neurodegeneration. Neural Plast 2019, 2019:4209475.

99. Lauretti E, Dincer O, Praticò D: Glycogen synthase kinase-3 signaling in Alzheimer’s disease. Biochim Biophys Acta Mol Cell Res 2020, 1867(5):118664.

100. Zhong MZ, Peng T, Duarte ML, Wang M, Cai D: Updates on mouse models of Alzheimer’s disease. Mol Neurodegener 2024, 19(1):23.

101. Mol P, Gopalakrishnan L, Chatterjee O, Mangalaparthi KK, Kumar M, Durgad SS, Nair B, Shankar SK, Mahadevan A, Prasad TSK: Proteomic Analysis of Adult Human Hippocampal Subfields Demonstrates Regional Heterogeneity in the Protein Expression. J Proteome Res 2022, 21(10):2293–2310.

102. Mendis LH, Grey AC, Faull RL, Curtis MA: Hippocampal lipid differences in Alzheimer’s disease: a human brain study using matrix-assisted laser desorption/ionization- imaging mass spectrometry. Brain Behav 2016, 6(10):e00517.

103. Deuker L, Doeller CF, Fell J, Axmacher N: Human neuroimaging studies on the hippocampal CA3 region - integrating evidence for pattern separation and completion. Front Cell Neurosci 2014, 8:64.

104. Tzakis N, Holahan MR: Social Memory and the Role of the Hippocampal CA2 Region. Front Behav Neurosci 2019, 13:233.

105. Bartsch T, Döhring J, Rohr A, Jansen O, Deuschl G: CA1 neurons in the human hippocampus are critical for autobiographical memory, mental time travel, and autonoetic consciousness. Proc Natl Acad Sci U S A 2011, 108(42):17562–17567.

106. Gilbert PE, Kesner RP, Lee I: Dissociating hippocampal subregions: double dissociation between dentate gyrus and CA1. Hippocampus 2001, 11(6):626–636.

