## Supplementary file 1 for "Mapping the spatial proteomic signature of dorsal and ventral hippocampus in a mouse model of early Alzheimer’s disease: changes in synaptic plasticity-related proteins associated with sexual dimorphism"

### Proteins validation by western blot

Western blot analyses were conducted with two selected altered proteins to validate expression changes found by MALDI imaging analyses.

RCAN1 was down-regulated according to MALDI imaging analysis and western blot revealed a significant treatment effect ($F_{(1,28)}$ = 9.776, $p$ = 0.0041). Although no difference between males and females was observed ($F_{(1,28)}$ = 0.4843, $p$ = 0.4922), *post hoc* analysis revealed that the effect was specifically in males oA$\beta_{1\text{-}42}$ treated mice (Figure S1A).

On the other hand, GluR5 was found upregulated by MALDI imaging analysis as well as by western blot (treatment effect: $F_{(1,20)}$ = 5.721, $p$ = 0.0267). However, *post hoc* analysis did not indicate to which specific group the effect was due. Once again, no sex difference was found ($F_{(1,20)}$ = 0.2243, $p$ = 0.6409; Figure S1B).

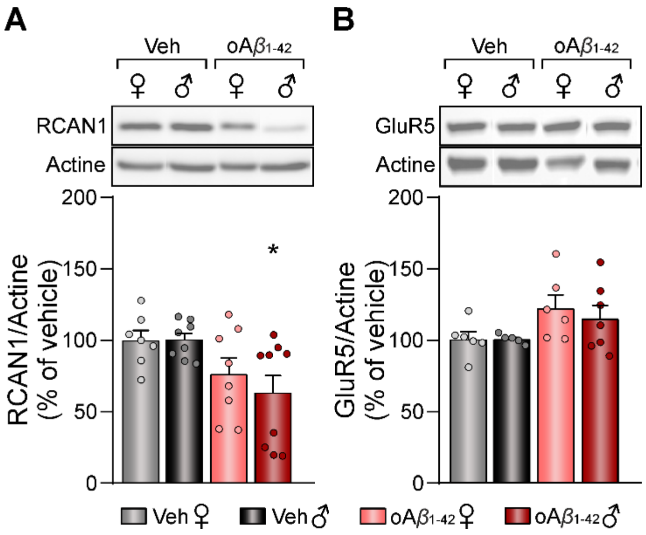


**Figure S1. Validation of hippocampal proteins by western blot.** Relative expression of RCAN1 **(A)** or GluR5 **(B)** in vehicle and oA$\beta_{1\text{-}42}$ treated mice and representative western blots. Data is expressed as mean ± SEM of the target protein/actine as a loading control, and as percentage (%) of the control (vehicle) group of the corresponding sex.

N vehicles: males = 5-8 and females = 6-7; N $oA\beta_{1-42}$: males = 7-9 and females = 6-8. $oA\beta_{1-42}$, Amyloid-$\beta_{1-42}$ oligomers; veh, vehicle. * $p < 0.05$ *vs.* vehicle of the corresponding sex.
